# Detecting and visualising the impact of heterogeneous evolutionary processes on phylogenetic estimates

**DOI:** 10.1101/828996

**Authors:** Lars S Jermiin, David R Lovell, Bernhard Misof, Peter G Foster, John Robinson

**Affiliations:** Land & Water, CSIRO, Acton, ACT 2601, Australia; Research School of Biology, Australian National University, Acton, ACT 2601, Australia; School of Biology & Environmental Science, University College Dublin, Belfield, Dublin 4, Ireland; Earth Institute, University College Dublin, Belfield, Dublin 4, Ireland; Data61, CSIRO, Acton, ACT 2601, Australia; School of Electrical Engineering & Computer Science, Queensland University of Technology, Brisbane, QLD 4001, Australia; Zoologisches Forschungsmuseum Alexander Koenig, 53113 Bonn, Germany; Department of Life Sciences, Natural History Museum, London, SW7 5BD, UK; School of Mathematics & Statistics, University Sydney, Sydney, NSW 2006, Australia

**Author notes:** Correspondence: Lars S Jermiin, Research School of Biology, Australian National University, Canberra, ACT, 0200, Australia;.

**Keywords:** Evolution under stationary conditions, Matched-pairs test of symmetry, PP plot, Heat map, Historical signal, compositional signal, compositional distance, networks

## Abstract

Most model-based molecular phylogenetic methods assume that the sequences diverged on a tree under homogeneous conditions. If evolution occurred under these conditions, then it is unlikely that the sequences would become compositionally heterogeneous. Conversely, if the sequences are compositionally heterogeneous, then it is unlikely that they have evolved under homogeneous conditions. We present methods to detect and analyse heterogeneous evolution in aligned sequence data and to examine—visually and numerically—its effect on phylogenetic estimates. The methods are implemented in three programs, allowing users to better examine under what conditions their phylogenetic data may have evolved.

---

Most model-based molecular phylogenetic methods assume that the sequences of nucleotides or amino acids have evolved along the edges of a single bifurcating tree. Often, the methods also assume that the evolutionary processes operating at the variable sites of these data (i.e., the sites that are free to evolve) can be approximated by independent and identically-distributed (*iid*) Markovian processes. Furthermore, it is often assumed that the evolutionary processes were stationary, reversible and homogeneous (SRH) (for details, see Bryant et al. 2005; Jayaswal et al. 2005; Ababneh et al. 2006a,b; Jermiin et al. 2017), with the term homogeneity implying time-homogeneity (i.e., a constant rate of change between two points in time).

In practice, when DNA has evolved under these conditions, commonly-used phylogenetic methods are likely to identify the correct topology (Huelsenbeck and Hillis 1993; Hillis et al. 1994a,b). However, the same methods may not be capable of identifying the correct topology when DNA has evolved under more complex conditions (Huelsenbeck and Hillis 1993; Hillis et al. 1994a,b; Ho and Jermiin 2004; Jermiin et al. 2004). One reason for this failure is that the strength of the *historical signal* (i.e., the signal in DNA that is due to the order and time of divergence events) decays over time (Ho and Jermiin 2004) whereas the strength of the *non-historical signals* (Grundy and Naylor 1999) may increase over time (Fig. 1). This may lead to situations, where the non-historical signals—individually or jointly—may become stronger than the historical signal (Ho and Jermiin 2004). Unless phylogenetic methods are able to distinguish historical signals from non-historical signals, the latter may be misinterpreted as being part of the historical signal. This is because the non-historical signals are also *phylogenetic signals*.

**Figure 1:**
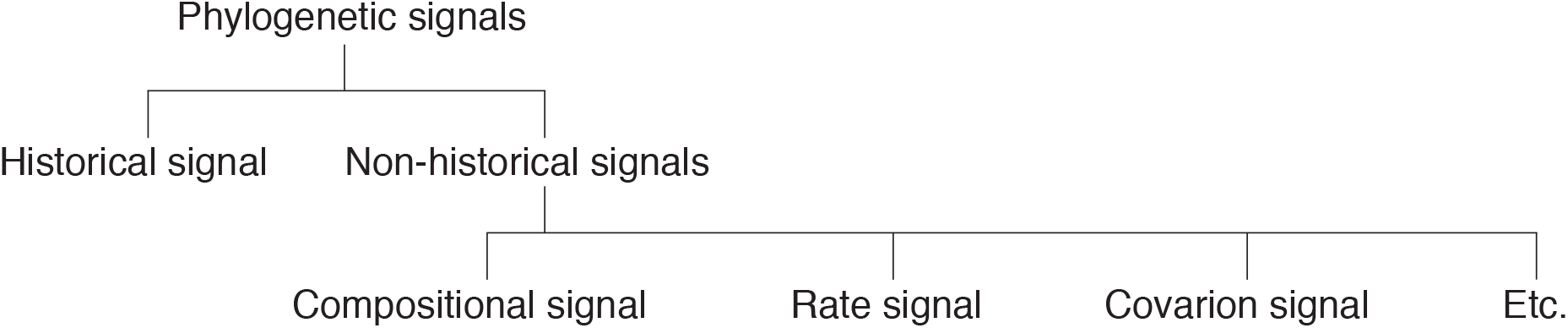
The phylogenetic signals (i.e., signals in phylogenetic data that, on their own, can generate a phylogeny), partitioned into some of its constituent components. Phylogenetic studies often aim to extract a historical signal from phylogenetic data. However, the accuracy of these studies depends not only on how decayed the historical signal is (Ho and Jermiin 2004) but also on whether non-historical signals have arisen over the course of time. The non-historical signals include the compositional signal (caused by non-homogeneous site patterns in the data), the rate signal (caused by independently evolving sites evolving at different rates), the covarion signal (caused by sites not evolving independently). Non-historical signals may bias phylogenetic estimates unless properly accounted for.

The non-historical signal is a mixed bag of signals that may arise over time due to temporal variations in site- and lineage-specific evolutionary processes. For example, when the homologous sites in a pair of sequences evolve under different conditions, evolutionary processes cannot be homogeneous, and compositional heterogeneity across the sequences may arise. When this happens, there is a *compositional signal* in the data (Fig. 2). On the other hand, when compositional heterogeneity is found across an alignment of homologous sequences, there is evidence of evolution under non-stationary conditions.

**Figure 2:**
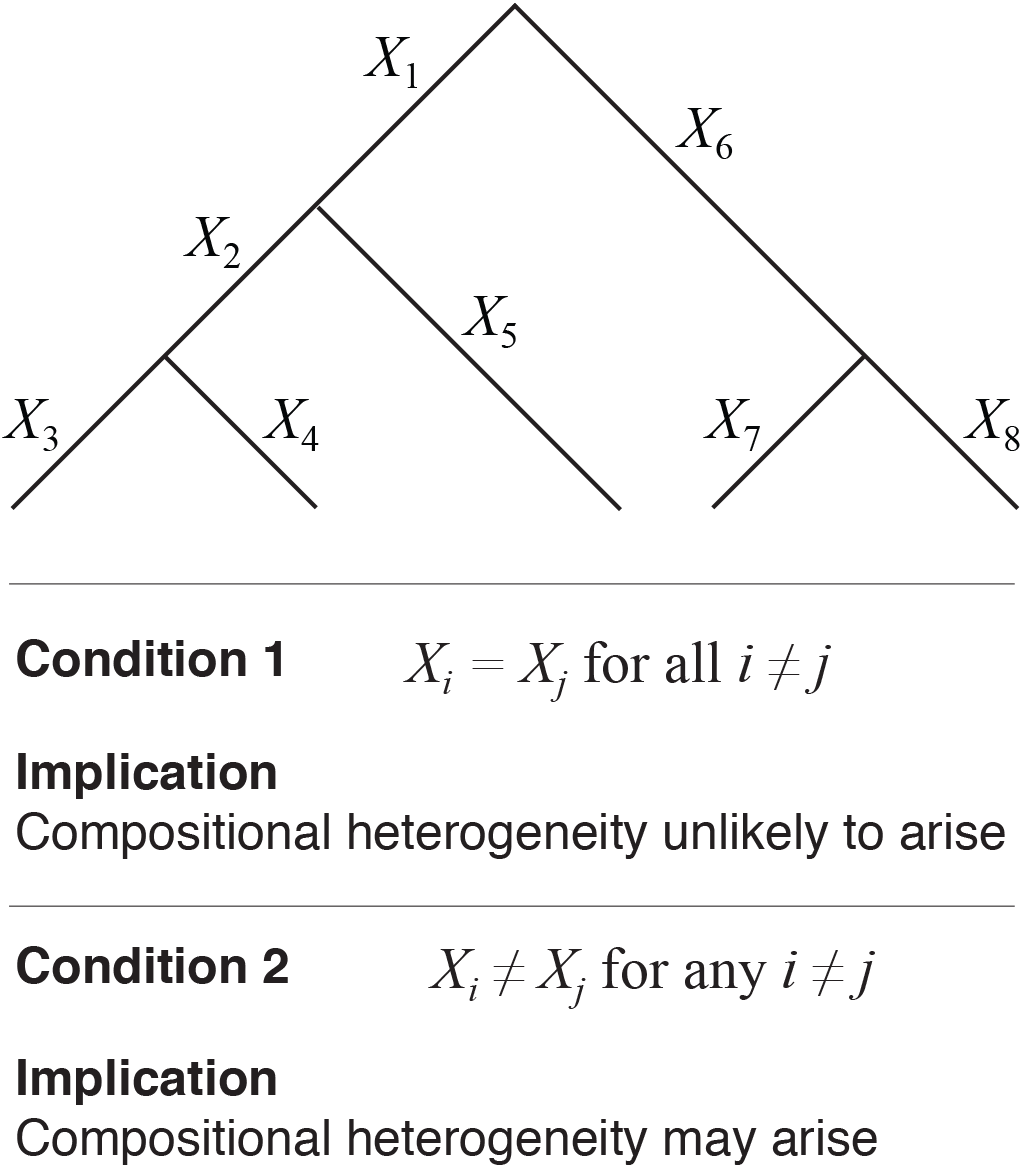
The phylogenetic challenge, illustrated using a nucleotide sequence evolving over a rooted 5-tipped tree with eight Markovian processes (i.e., *X*_1_, …, *X*_8_) distributed over the edges. Each site in the sequence evolving over this tree is governed by these eight edge-specific Markov processes. If *X*_*i*_ = *X*_*j*_ for all *i ≠ j*, compositional heterogeneity across the descendant sequences is unlikely to arise. Otherwise, it may arise.

Several methods have been developed to detect compositional heterogeneity across homologous sequences (reviewed in Jermiin et al. (2004, 2009)), but doubt remains about what method is most appropriate (cf. Jermiin et al. (2004) and Duchêne et al. (2017)). To resolve this matter and to empower concerned users of phylogenetic methods, we present software to detect and visualise compositional heterogeneity across aligned sequence data. The software also facilitates assessment of the impact of compositional heterogeneity on inferred phylogenetic trees and networks.

## Methodology

### Background

Consider a nucleotide sequence that evolves over the edges of a rooted tree (Fig. 3), and assume that the 90 sites in this sequence evolve under *iid* conditions. At time *t*_0_, the ancestral sequence, `Seq0`, evolves along an ancestral edge in the tree (Fig. 3a). At time *t*_1_, the sequence meets a bifurcation in the tree, and it becomes two identical sequences, `Seq1` and `Seq2` (Fig. 3b). At time *t*_2_, the two sequences have evolved further under independent evolutionary processes (Fig. 3c), so they are unlikely to be the same. The sequences at *t*_0_, *t*_1_ and *t*_2_ are shown in Figure 3d.

**Figure 3:**
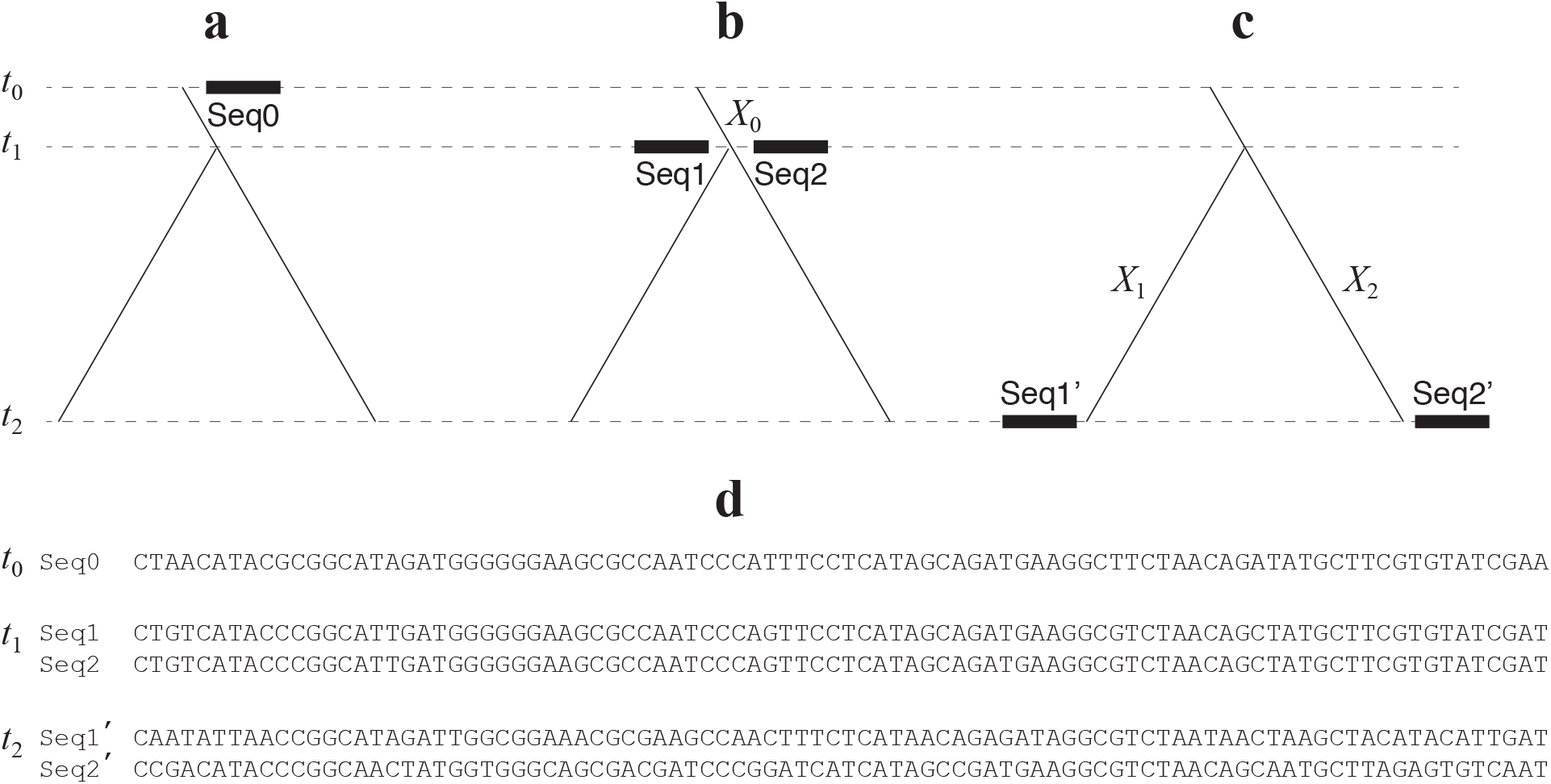
Rooted phylogenetic tree with the ancestral sequence evolving along the root edge (**a**) and, later on, at the start (**b**) and the end (**c**) of the bifurcation. The evolutionary processes operating over the three edges are marked *X*_0_, *X*_1_ and *X*_2_. The corresponding sequences from the three points in time (i.e., *t*_0_, *t*_1_ and *t*_2_) are shown in panel **d**.

Methodologically, the challenge now is to extract as much information as possible from the alignment of `Seq1′` and `Seq2′` (e.g., to infer the time elapsed since the bifurcation at *t*_1_). One way to extract information from such a data set is to consider the ratio of the number of sites where the sequences differ to the total number of sites compared. This yields a metric called the *p* distance (the *p* distance between `Seq1′` and `Seq2′` is 42/90). Another way to do this is to use a divergence matrix (**N**). For `Seq1` and `Seq2`, we get:

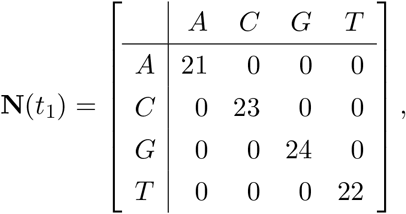

while for `Seq1′` and `Seq2′` we get:

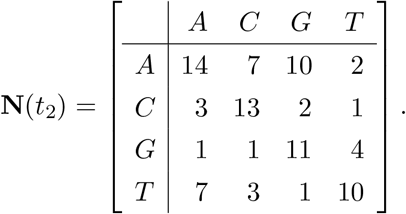

The only difference between **N** and the alignments in Figure 3d is that information about the order of sites in the alignment is lost in the divergence matrix. However, as these sites are assumed to have evolved independently, this loss of information is of no consequence for most commonly-used phylogenetic methods.

Given **N**, we can obtain the *p* distance or any other evolutionary distance, like the F81 distance (Felsenstein 1981). Likewise, we can determine whether two sequences have diverged under homogeneous conditions. If the distributions of *X*_1_ and *X*_2_ are equal, then the sequences will have evolved under homogeneous conditions. Assuming evolution under homogeneous conditions, the divergence matrix should be approximately symmetrical (i.e., if **N** = {*n*_*ij*_}, then *E*(*n*_*ij*_) = *E*(*n*_*ji*_) ∀ *i, j*; *E* denotes the expected value).

### The Matched-pairs Test of Symmetry

The matched-pairs test of symmetry is suitable for testing whether *E*(*n*_*ij*_) = *E*(*n*_*ji*_). It is computed using:

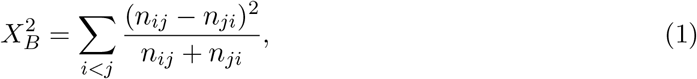

which, assuming homogeneous conditions, is asymptotically distributed as a *χ*^2^ variate on *ν* = *c* × (*c* − 1)/2 degrees of freedom, where *c* denotes the number of unique letters in the sequences’ alphabet (for DNA, *c* = 4). Given 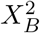 and *ν*, it is easy to obtain the probability of getting a test statistic that equals or exceeds 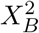, given *ν* (i.e., 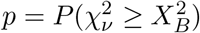). In this regard, it is worth remembering that 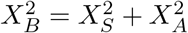, where 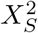 is the test statistic from the matched-pairs test of marginal symmetry (Stuart 1955) while 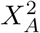 is the test statistic from the matched-pairs test of internal symmetry (Ababneh et al. 2006b). It is also worth pointing out that if for any of the comparisons *n*_*ij*_ + *n*_*ji*_ = 0, the entry is ignored and *ν* is reduced by 1.

The matched-pairs test of symmetry was devised by Bowker (1948) and introduced to molecular phylogenetics by Tavaré (1986). Subsequent attempts to promote this test as the best approach to test homogeneity of the evolutionary processes (Lanave and Pesole 1993; Waddell and Steel 1997; Waddell et al. 1999; Ababneh et al. 2006b) were largely unsuccessful, with one opponent stating that the test “is hardly necessary because typical phylogenetic datasets are large and can reject the null hypothesis with ease” (Yang 2014). That is an odd statement, as it recommends ignoring a reason for systematic error. More recently, Duchêne et al. (2017) used a test described by Foster (2004), which tests the fit of the compositional component of the (stationary, in this case) model to the data. This test uses a contingency table made up of *c* marginal sums. However, unlike the standard *r × c* contingency table test of homogeneity, the test statistic is not compared to the *χ*^2^ distribution but to a simulated null distribution obtained on the basis of the tree and the (possibly non-stationary) model of evolution being tested. In other words, it is a test of model fit—it needs to be used after the tree and model of evolution have been specified. Thus, it is akin to the Goldman-Cox test of goodness-of-fit (Goldman 1993), which uses simulations to assess the significance of a statistic.

While the test used by Foster (2004) tests marginal compositions, it ignores the homology statements that alignments represent. The impact of doing so can be dramatic, as the following example reveals. The three divergence matrices, left to right, are the products of increasingly dissimilar evolutionary processes:

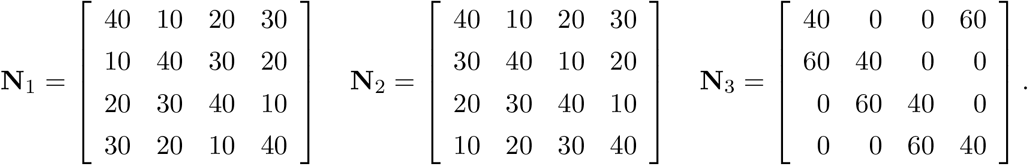

In the first case (**N**_1_), there is no evidence that the evolutionary processes might have been different, while in the other cases (**N**_2_ and **N**_3_), the evidence of that is clearer. However, it is also clear that the three matrices have the same marginal distribution, so Foster’s (2004) test cannot detect this type of lineage-specific heterogeneity in the evolutionary processes. Foster’s (2004) test is similar to Stuart’s (1955) matched-pairs test of marginal symmetry. If the aim is to test the fit between tree, model and data, then it would be appropriate to use Foster’s (2004) test or the Goldman-Cox test of goodness of fit (Goldman 1993). On the other hand, if the aim is to test whether sequences are consistent with the assumption of evolution under stationary conditions, then Stuart’s (1955) matched-pairs test of marginal symmetry is recommended (Ababneh et al. 2006a). Stuart’s (1955) matched-pairs test of marginal symmetry, like Bowker’s (1948) matched-pairs test of symmetry, assumes aligned data but not a tree or model, so it is useful for screening phylogenetic data *before* they are analysed. On the other hand, Foster’s (2004) test is applicable *after* this analysis, can be used with non-stationary models, and is not restricted to sequence pairs.

### The PP Plot

If we wish to apply the matched-pairs test of symmetry to an alignment with more than two sequences, then the problem of multiple comparisons arises. For example, if a data set contains 22 sequences, then there will be 22 × 21/2 = 231 *p*-values to interpret, one for each pair of sequences. However, the *p*-values are not independent, so they must be interpreted jointly. This can be done using a PP-plot, which displays observed *p*-values against expected *p*-values. If evolution occurred under homogeneous conditions, then the 231 *p*-values will be distributed as a uniform random variable on (0,1). Given this expectation, we can evaluate whether the data set, as a whole, meets the assumption of evolution under homogeneous conditions.

To demonstrate the merits of the PP plot, we analysed an alignment of simulated nucleotides generated under time-reversible conditions on a 22-tipped tree (Fig. 4a). The PP plot in Figure 4b shows the result from data generated under the null hypothesis. As expected, the 231 dots are distributed along the diagonal, with ∼ 5% of them (12) below 0.05 (i.e., the horizontal line in Fig. 4b). None of the observed *p*-values fell below the 5% family-wise error rate (i.e., 0.05/231 = 0.000216). The PP plot shows the distribution to expect when the data have evolved under homogeneous conditions. This interpretation is consistent with those in Schweder and Spjøtvoll (1982) and Vera-Ruiz et al. (2014).

**Figure 4:**
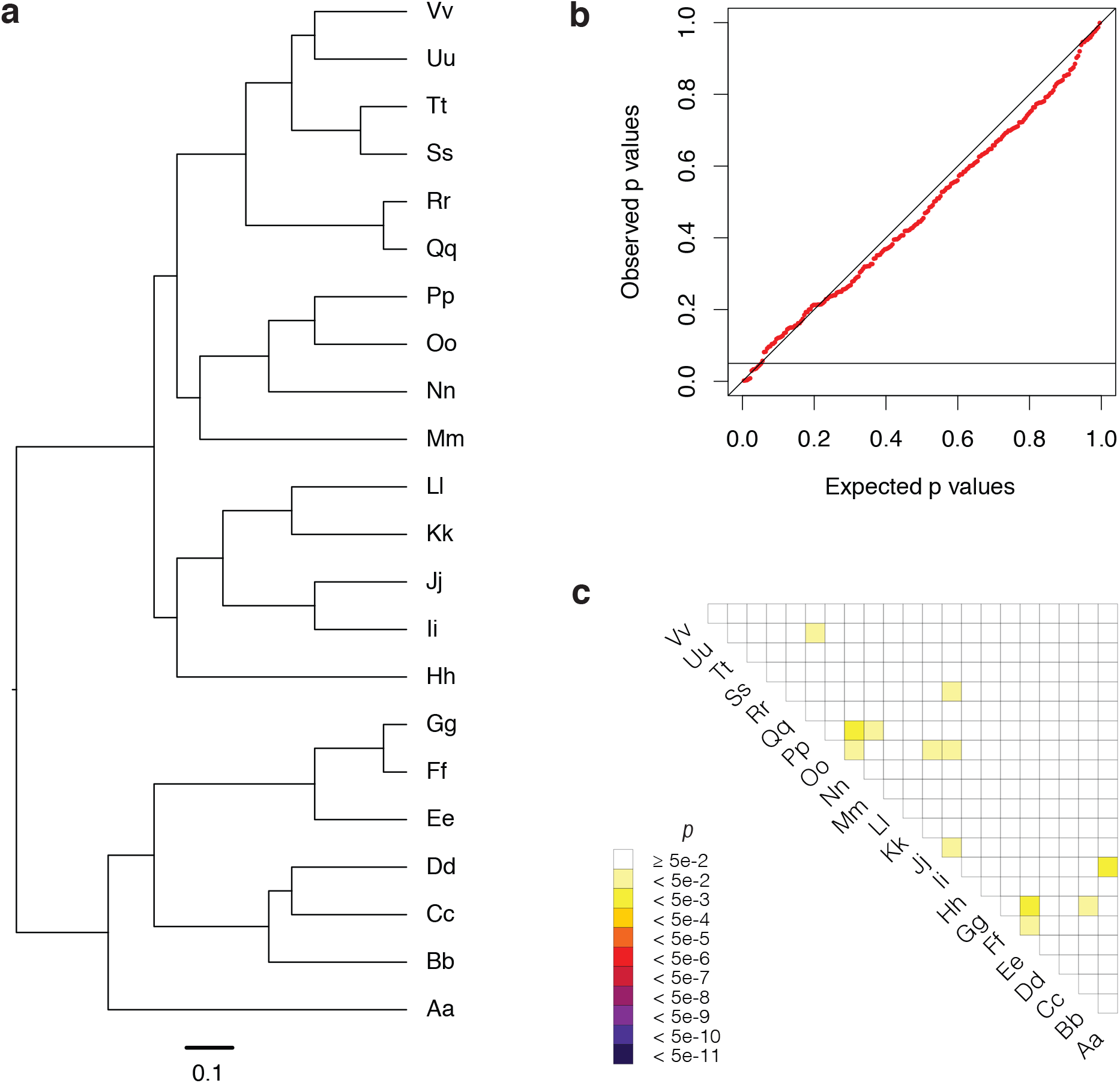
Using Seq-Gen (Rambaut and Grassley 1997), nucleotide sequences containing 100,000 sites were generated by simulation on a 22-tipped tree (**a**) under the GTR (Tavaré 1986) model of sequence evolution with the following parameters: *S* = [0.8, 0.4, 0.2, 0.1, 0.05, 0.025], *π* = [0.4, 0.3, 0.2, 0.1], *pI* = 0.15, and a continuous Γ distribution with *α* = 2.7. The resulting sequences were then analysed using the matched-pairs test of symmetry. The resulting 231 *p*-values were finally presented in a PP plot (**b**) and in a heat map (**c**).

### The Heat Map

A PP-plot that deviates noticeably from that shown in Fig. 4b (e.g., the dots are not distributed along the diagonal; more than 5% of the observed *p*-values are below 0.05; the smallest observed *p*-value is below a 5% family-wise error rate), suggests that some of the sequences have evolved under heterogeneous conditions. However, the PP plot cannot identify the ‘offending’ sequences, but a color-coded heat map with the observed *p*-values can. Figure 4c shows the heat map corresponding to the data in Figure 4b. Each pixel is color-coded according to the *p*-value for the corresponding pair of sequences. Most of the pixels are white because the *p*-values are ≥ 0.05. Some pixels are yellow, but none of them are darker; this is consistent with the condition under which the sequences were generated.

When a heat map differs noticeably from that in Figure 4c, it allows us to identify sequences that are unlikely to have evolved under the same conditions. For example, if all but one of the sequences evolved under homogeneous conditions, then that would result in a heat map where a row and/or column has darker pixels. The color of a pixel depends on the probability that the corresponding pair of sequences have evolved under homogeneous conditions. A dark row and/or column identifies an offending sequence, which then can be removed if it is insignificant to the phylogenetic question. Figure 6 of Jayaswal et al. (2014) shows such a heat map (in this case the offending sequences could not be removed).

When two or more sequences are regarded as offending, we might ask whether the data can be grouped into subsets of sequences that are consistent with evolution under homogeneous conditions. To do so, one simply needs to permute the rows and columns of the heat map, or reorder the sequences in the alignment before analysing the data again. Figures 6 and 7 of Jermiin et al. (2017) show two heat maps for the same data, obtained before and after a permutation of the rows and columns of the heat map. In the first figure, several small sets of sequences appear to have evolved under homogeneous conditions. However, the second figure reveals that many of these subsets can be merged into larger subsets of sequences that appear to have evolved under different homogeneous conditions. In summary, the PP plot and heat map provide researchers an opportunity to survey their data far more thoroughly before model selection and phylogenetic analysis.

### Compositional Distances

A compositional signal may arise when sequences diverge under non-homogeneous conditions. If such a signal emerges, its amplitude can be measured using distance metrics that quantify departure from symmetry of a divergence matrix. Compositional distances are appropriate for vectors of non-negative values that carry information in their relative (not absolute) amounts (Aitchison 1986; Egozcue and Pawlowsky-Glahn 2011), like those in a divergence matrix. Compositional distances may be used to infer trees and networks, revealing relationships based solely on compositional differences. These trees and networks may uncover a compositional signal’s potential impact on phylogenetic estimates.

Given **N** (for nucleotides):

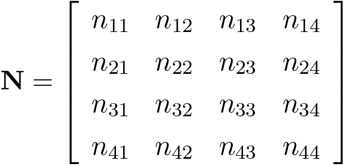

we can define two vectors that relate to the off-diagonal elements of the upper and lower triangles:

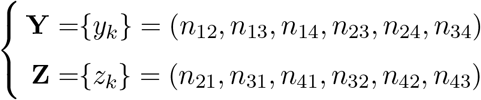

Given **Y** and **Z** for a *c*-state alphabet (e.g., *c* = 20 for protein), it is possible to compute three compositional distances:

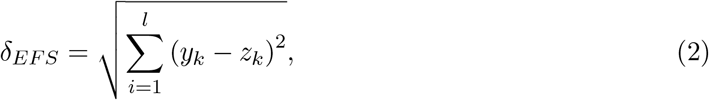

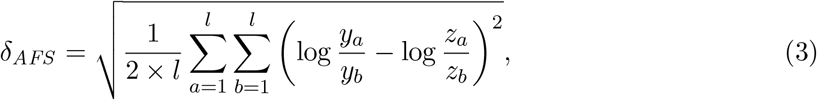

and

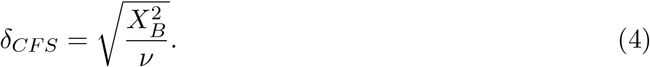

Here, *δ*_*EFS*_, *δ*_*AFS*_, and *δ*_*CFS*_ respectively denote the Euclidean distance, Aitchison’s (1986) distance, and a distance metric closely related to Bowker’s (1948) matched-pairs test of symmetry, and *l* is the number of elements in **Y** and **Z**. The Euclidean distance measures the distance between two points in Euclidean space, taking no account of sign or scale, so they are not appropriate for count data. One more appropriate metric is that of Aitchison (1986); for a comparison of these distance metrics, see Lovell et al. (2011). One undesirable property of *δ*_*AFS*_ is that it is zero when *n*_*ij*_/*n*_*ji*_ is constant, and will be small if this is even approximately so. Because of this, Aitchison’s (1986) distance is not suitable for data used to measure lack of symmetry in divergence matrices. Instead, we may use *δ*_*CFS*_, which has the advantage of being able to accommodate that comparisons between different pairs of sequences may be associated with different degrees of freedom (*ν*). Note that *δ*_*CFS*_ ≥ 0.0, and that *δ*_*CFS*_ is not an evolutionary distance in the sense that the LogDet (Lockhart et al. 1994; Steel 1994) or paralinear (Lake 1994) distances are.

### The Nature of Bias in Phylogenetic Estimates

It is difficult to detect bias in phylogenetic estimates from real sequence data, but it is well known that bias may manifest itself in at least two ways:

1. The topology of the tree (or network) is affected, implying that the length of at least some of the edges (or weights of some of the splits; ‘weight’ is analogous with length, in the sense of Huson and Bryant (2006)) also will be affected, or
2. The topology is unaffected but the length of the edges in the tree (or the weights of the splits in the network) may be affected.

Both of these biases are cause for concern, even if only the topology is of interest, because the topology is a discrete entity, whose accuracy often is dependent on the accuracy of the estimates of the other parameters. The challenge is to get all the estimates as accurate as possible without increasing the variance or the bias of these estimates (Dziak et al. 2019). In other words, both over- and under-parameterisation of the data should be avoided.

### Visualising the Effect of Compositional Heterogeneity on Trees and Networks

Given a distance matrix **D**_*CFS*_ with estimates of *δ*_*CFS*_, we may infer a *compositional tree, 𝒯*, and a *compositional network, 𝒩*. This can be done by using programs like FastME (Lefort et al. 2015) and SplitsTree4 (Huson and Bryant 2006). Such structures display the relationships among sequences based solely on compositional distances, so they should not be interpreted as if they were phylogenetic trees or phylogenetic networks. Sequences that are compositionally similar may not be close in an evolutionary sense, and sequences that are compositionally dissimilar may not be distantly-related in an evolutionary sense. The advantage of using data-display networks to reveal conflicting signals in phylogenetic data has already been demonstrated by Morrison (2010), so it will not be reiterated here.

Consider a data set that has been found to violate the phylogenetic assumption of evolution under homogeneous conditions. In such a case, one might wish to know whether the compositional signal has become so strong that it might bias a phylogenetic estimate, unless it is properly accounted for.

To demonstrate the benefit of using *𝒯* and *𝒩*, we analysed an alignment of five 16S rRNA sequences from bacteria, first analysed phylogenetically by Embley et al. (1993) and then by Galtier and Gouy (1995), Mooers and Holmes (2000), Foster (2004), and Jayaswal et al. (2005, 2007). For these data, **D**_*EFS*_, **D**_*AFS*_ and **D**_*CFS*_ are

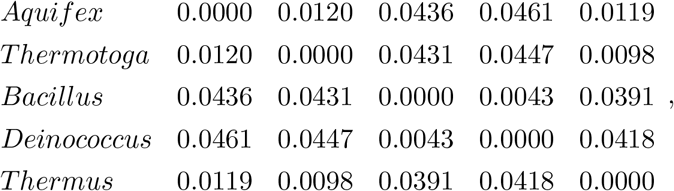

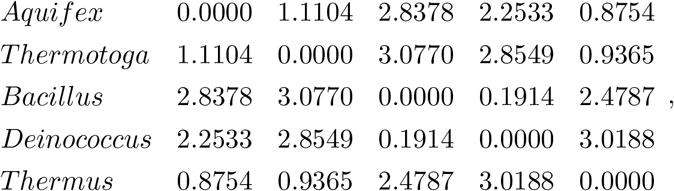

and

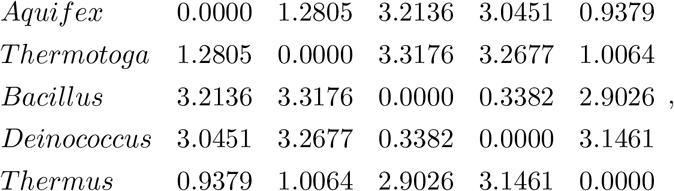

respectively (the values in **D**_*EFS*_ and **D**_*AFS*_ were obtained using Homo v1.3: Rouse et al. 2013). The three matrices differ, reflecting the differences between Equations 2, 3 and 4. Interestingly, the elements of **D**_*EFS*_, **D**_*AFS*_ and **D**_*CFS*_ appear to be highly correlated (i.e., carrying quite similar information), but this is not always the case (e.g., if **Y** *∝* **Z**).

Figures 5a and 5b shows a BioNJ tree (Gascuel 1997) and a Neighbor-Net (Bryant and Moulton 2004), both inferred from **D**_*CFS*_ using SplitsTree4 (Huson and Bryant 2006). The compositional tree (*𝒯*) has a long internal edge (marked † in Fig. 5a) that separates *Deinococcus* and *Bacillus* from the other three species. The same appears to be the case for the compositional network (*𝒩*) in Fig. 5b. Indeed, *𝒩* is very *treelike*, because the split marked † in Figure 5b is 18.6 times longer than the second-longest alternative (marked ‡). In other words, *𝒯* and *𝒩* corroborate what is already known about these five sequences: *Deinococcus* and *Bacillus* are compositionally distinct from the other three species (Galtier and Gouy 1995; Jayaswal et al. 2005). However, in many other studies, such knowledge is not available or heeded. This is where compositional trees or networks become useful; not only do the topologies of *𝒯* and *𝒩* identify the compositionally most similar sequences, they also reveal where the biggest differences are—and as compositional differences grow, so do the length of edges in *𝒯* and splits in *𝒩*. Importantly, compositional networks are able to reveal conflicting information in multiple sequence alignments that compositional trees cannot reveal (because the latter are constrained to be acyclic graphs: Penny et al. 1992). Therefore, during the exploratory phase of assessing compositional heterogeneity, using *𝒩* may be better than using *𝒯*.

**Figure 5:**
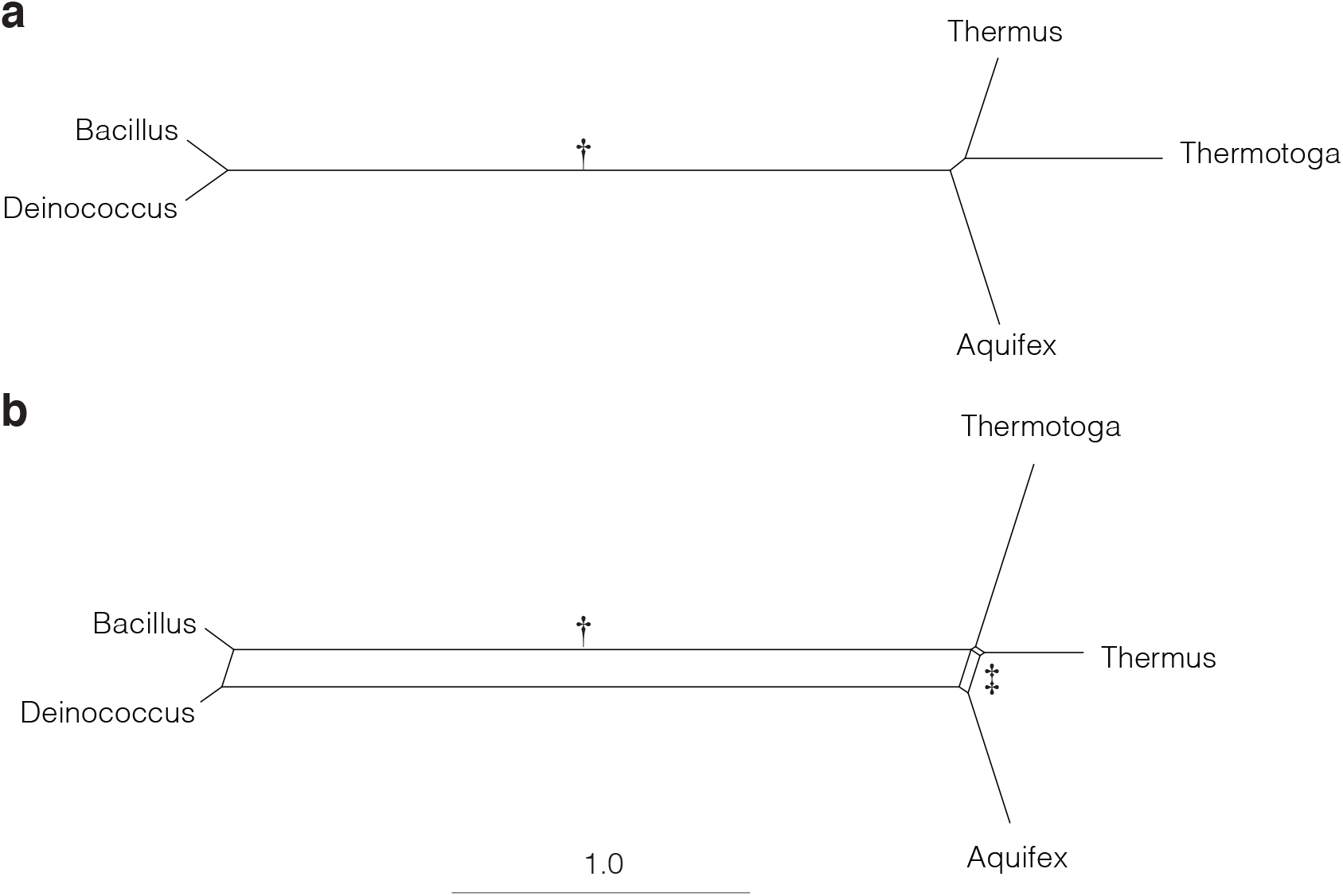
A compositional tree (**a**) and a compositional network (**b**), inferred from a matrix of compositional distances (**D**_*CFS*_) obtained from an alignment of bacterial 16S rRNA sequences. The tree and the network are drawn to scale. The characters † and ‡ point to splits that are referred to in the text.

**Figure 6:**
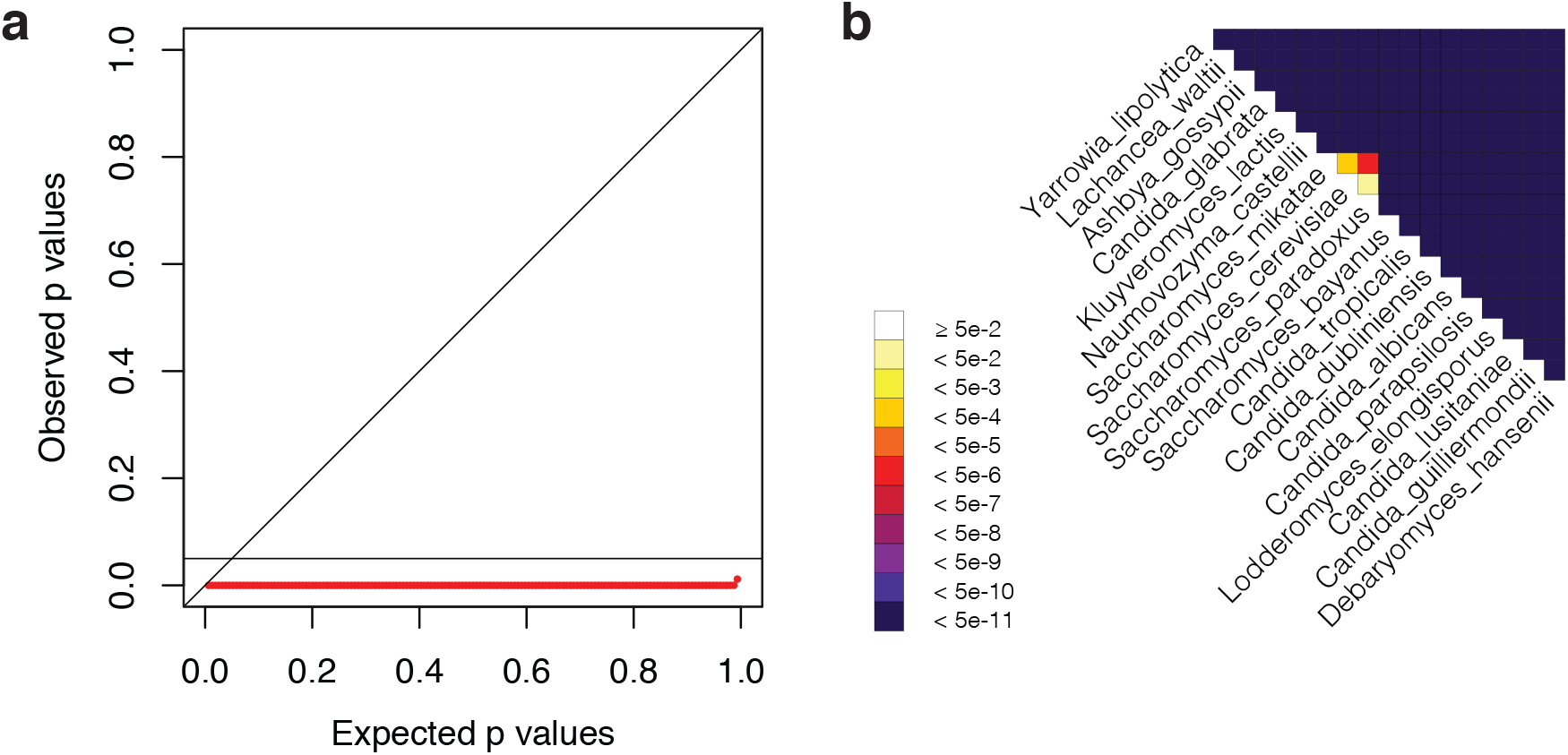
Visual output from our study of the alignment of amino acids from Butler et al. (2009). (**a**) PP plot showing that the data set, as a whole, is unlikely to have evolved under homogeneous conditions. (**b**) Heat map identifying the least offending sequences (*Saccharomyces cereviciae* and *S. paradoxus*). In Butler et al. (2009), *Lachancea waltii* was called *Kluveromyces waltii* and *Naumovozyma castelliii* was called *S. castellii*.

**Figure 7:**
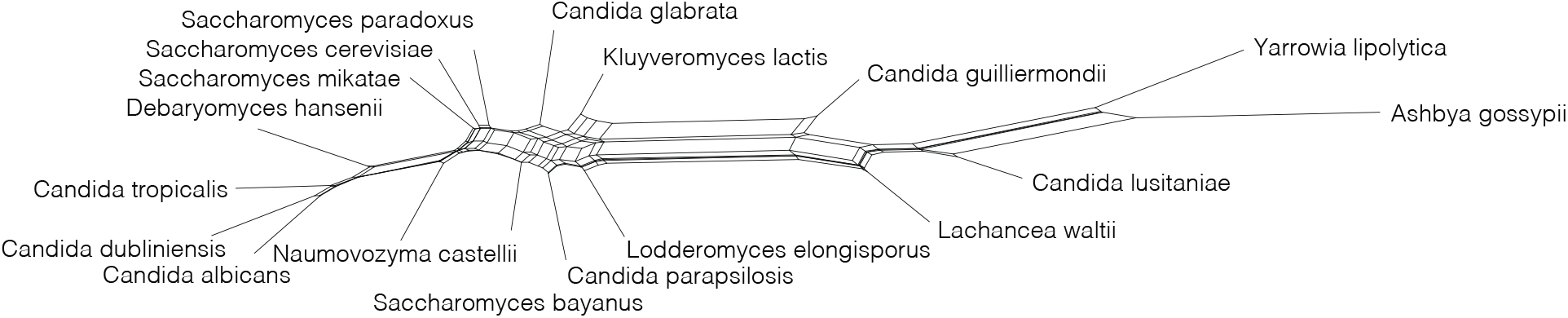
A compositional network inferred by SplitsTree4 (Huson and Bryant 2006) from a matrix of compositional distances (**D**_*CFS*_) obtained from an alignment of amino acids by Butler et al. (2009). In Butler et al. (2009), *Lachancea waltii* was called *Kluveromyces waltii* and *Naumovozyma castelliii* was called *S. castellii*.

### Congruence between Phylogenetic and Compositional Trees

Interestingly, the split observed between *Deinococcus* and *Bacillus* and the other three species (Fig. 5) is also found in optimal phylogenetic trees inferred under different time-reversible Markovian models of sequence evolution (Jayaswal et al. 2005, 2007). At least two explanations may be given for this congruence of splits:

1. The historical and compositional signals in the data are aligned, implying that the historical signal is augmented by the compositional signal. A consequence of this is that the inferred topology may be correct. However, estimates of edge lengths may still be biased; this could lead to bias in estimates of divergence dates.
2. The historical and compositional signals are not aligned, implying that the historical signal might be undermined by the compositional signal. This would entail that the phylogenetic methods, unless specifically designed to accommodate a compositional signal, might misinterpret the compositional signal, as if it were the historical signal, and return a phylogenetic estimate with biases in both topology and edge lengths.

In the first explanation, the compositional signal may be stronger than the historical signal but because the two signals are aligned, this has no adverse effect on the inferred topology; on the contrary, it may help us to identify the correct topology. In the second explanation, both the strength and the complexity of the compositional signal are likely to contribute to bias in phylogenetic estimates. Importantly, the identities and lengths of internal edges in the true tree are both factors contributing to the success or failure of phylogenetic inference (Jermiin et al. 2004), but neither of these factors is known (except for in simulation-based studies).

The problem with these two explanations is that they apply equally well to many studies of compositionally heterogeneous phylogenetic data sets and that we do not know which one is right. It is not wise to argue that other phylogenetic estimates corroborate a current phylogenetic hypothesis, unless bias due to model misspecification has been ruled out for *all the data sets* being compared. In the present case, the matter was resolved by analysing the alignment using a model that was heterogeneous over the tree (Foster 2004) and by using the general Markov model of sequence evolution (Jayaswal et al. 2007). However, this is rarely done.

### Testing for Similarity between Phylogenetic and Compositional Trees

Often phylogenetic data contain more than five sequences and it may be less clear (than e.g., Fig. 5) whether a compositional signal contributed adversely to a phylogenetic estimate. In such cases, it may be useful to compare the phylogenetic tree (*𝒯*_*r*_ — inferred directly from the sequence alignment) and the compositional tree (*𝒯*_*c*_ — inferred from the corresponding matrix of compositional distances (**D**_*CFS*_)). In such instances, the distance between *𝒯*_*r*_ and *𝒯*_*c*_ must first be obtained. Reviewing the performance of tree-comparison metrics, Kuhner and Yamato (2015) found that Nye et al.’s (2006) metric, which is based on topology only, is superior for dissimilar trees. Their metric, *δ*_*Align*_, which measures how well two trees align to each other, was revealed to be better than four other tree-distance metrics, including the Robinson and Foulds (1981) metric and the Path Difference metric (Williams and Clifford 1971; Penny et al. 1982).

When comparing *𝒯*_*r*_ and *𝒯*_*c*_, a critical question is whether they are more similar, or dissimilar, to one another than random trees are to each other. If the evolutionary process of sequence data is modelled accurately, there is no reason to presume that *𝒯*_*r*_ and *𝒯*_*c*_ will be more similar, or dissimilar, to one another, than two random trees are. Thus, we may formulate a testable null hypothesis. H_0_: *𝒯*_*r*_ and *𝒯*_*c*_ are neither more similar, or dissimilar, to each other than random trees are.

To execute this test, we first calculate *δ*_*Align*_ for *𝒯*_*r*_ and *𝒯*_*c*_. Next, we generate, say, 2000 random trees and partition them into 1000 pairs. For each pair, we calculate 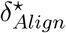, where the ‘star’ signals that this is an estimate obtained from random trees. Finally, the distribution of 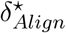 values is charted and the value of *δ*_*Align*_ for *𝒯*_*r*_ and *𝒯*_*c*_ is matched to this distribution. If the value of *δ*_*Align*_ falls well within the distribution of 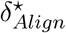, then the topologies of *𝒯*_*r*_ and *𝒯*_*c*_ are random with respect to each other; otherwise, they are more similar (e.g., if 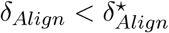 for all pairs) or dissimilar (e.g., if 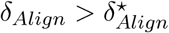 for all pairs) to each other than random trees are.

The method is illustrated in the biological example (below).

### Software

The methods described above are implemented in three programs.

#### Homo

Homo v2.0 is a complete re-development of previous versions of Homo (Rouse et al. 2013; http://www.csiro.au/Homo). Unlike the previous version, this one is written in C++ and designed for command line execution. Homo v2.0 includes corrections of errors found in the previous version, so Homo v1.3 should no longer be used. For each sequence pair, Homo executes the matched-pairs test of symmetry and returns:

- The probability (*p*) of getting the test statistic by chance (assuming evolution under homogeneous conditions),
- Euclidean distance (*δ*_*EFS*_) from full compositional symmetry of **N**,
- Euclidean distance (*δ*_*EMS*_) from marginal compositional symmetry of **N**,
- Our distance (*δ*_*CFS*_) from full compositional symmetry of **N**.

If any of the observed *p* values is below the 5% family-wise error rate, the program prints a warning to the user on the terminal. Homo is executed using the following commands:

~~~
homo <infile> <b|f> <1|…|31>
~~~

or

~~~
homo <infile> <b|f> <1|…|31> > README
~~~

where `infile` is a text file with an alignment of characters in the fasta format, `b|f` refers to whether a brief or full report of the results should be provided, and `1|…|31` refers the data type and how these data should be analysed. If `b` is used, Homo prints one line with key statistics to the user terminal; if `f` is used, it prints five files with the values of *p* and *δ*. A summary of the results is also be printed to the terminal.

Homo is designed to analyse alignments of nucleotides, di-nucleotides, codons, 10- and 14-state genotypes, and amino acids. If the `infile` contains sequences of:

- Single nucleotides (4-state alphabet), the sequences may be recoded into six 3-state alphabets or seven 2-state alphabets,
- Di-nucleotides (16-state alphabet; i.e., *AA, AC, …, TG, TT*), the sequences may be divided into alignments with 1st or 2nd position sequences,
- Codons (a 64-state alphabet; i.e., *AAA, AAC, …, TTG, TTT*), the sequences may be divided into three alignments with di-nucleotide sequences and three alignments with single-nucleotide sequences,
- Amino acids (a 20-state alphabet), the letters may be recoded to a 6-state alphabet. This type of recoding was recently used to study early evolution of animals (Feuda et al. 2017). Other types of recoding amino acids have been used (Kosiol et al. 2004; Susko and Roger 2007) but are not considered.

The 10- and 14-state genotype data cater for diploid and triploid genomes. For example, if a locus in a diploid genome contains nucleotides *A* and *G*, then the genotype sequence will contain an *R* at that locus. There are 10 distinguishable genotypes for each locus in diploid genomes and 14 for every locus in triploid genomes. For further detail about the data types and how the data may be analysed, simply type:

~~~
homo
~~~

on the command line and follow the instructions.

The output files from Homo fall into two categories: `.csv` files and `.dis` files. The `_Summary.csv` file contains all the estimates obtained for each pair of sequences. It can be opened and viewed by using, for example, Microsoft Excel. The `_Pvalues.csv` file contains all the *p* values set out in a format that can be read by HomoHeatMapper (see below). The three `.dis` files contain the *δ*_*CFS*_, *δ*_*EFS*_ and *δ*_*EMS*_ values, and can be analysed further using FastME (Lefort et al. 2015) and SplitsTree4 (Bryant and Moulton 2004).

#### HomoHeatMapper

HomoHeatMapper v1.0 is designed to generate a color-coded heat map from the `_Pvalues.csv` file. The colors used range from white (corresponding to *p* ≥ 0.05) to black (corresponding to *p* < 5 × 10^−11^). HomoHeatMapper is written in Perl and can be executed using the following command:

~~~
HomoHeatMapper -i <infile> -<t|f>
~~~

where `infile` must be the `_Pvalues.csv` file and where `t` and `f` stand for triangle and full, respectively. The output is an `.svg` file with a heat map in scalable vector graphics format. This file can be opened and processed using Adobe Illustrator.

#### RandTree

RandTree v1.0 is designed to generate random bifurcating trees from a set of labels. Starting from a rooted or unrooted tree with two or three tips, respectively, the tree is allowed to grow by randomly selecting tips, which will become bifurcating nodes in the tree. The probability that a tip is chosen equals 1/*n*, where *n* is the number of tips in the growing tree. Thus, the probability of selecting a given tip in a 16-leaf tree is 0.0625. Having obtained a random unlabelled tree, the labels are distributed randomly across the tips.

RandTree is a command-line tool written in C++. It is executed using:

~~~
randtree <infile> <r|u> <trees>
~~~

where `infile` is the text file with an unique taxon label on each line, `r|u` refers to whether the random trees should be rooted or unrooted, and `trees` refers to the number of random trees to generate. Trees generated by RandTree are printed in the Newick format to a text file, which can be used by other phylogenetic programs.

## Benchmarking

Recently, Naser-Khdour et al. (2020) applied the matched-pairs tests of symmetry (Bowker 1948), marginal symmetry (Stuart 1955), and internal symmetry (Ababneh et al. 2006b) to a panel of 35 published phylogenetic data sets with the aim to measure the prevalence and impact of model misspecification. Applying an implementation of these tests in IQ-TREE (Nguyen et al. 2015), their research revealed widespread evidence of evolution under non-SRH conditions, and that this appeared to impact the accuracy of phylogenetic estimates of these data inferred assuming evolution under SRH conditions. This observation complements that of a previous simulation-based study on the adverse impact of compositional heterogeneity on phylogenetic estimates (Jermiin et al. 2004).

We benchmarked Homo by comparing the result from the matched-pairs test of symmetry to those from the matched-pairs tests of symmetry, marginal symmetry, and internal symmetry, as implemented in TestSym (Ababneh et al. 2006b) and in IQ-TREE (Nguyen et al. 2015). In addition, we compared the result to that Foster’s (2004) test of homogeneity, as implemented in p4. We considered the alignment of `Seq1′` and `Seq2′` (Fig. 3d), and asked whether it is reasonable to assume that `Seq1′` and `Seq2′` diverged under homogeneous conditions (i.e., *X*_1_ = *X*_2_). The divergence matrix, **N**(*t*_2_), with its marginal frequencies, is reproduced here:

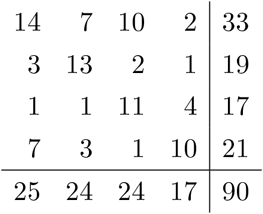

Table 1 shows the *p* values from different implementations of the matched-pairs tests of symmetry, marginal symmetry, and internal symmetry. As expected, Homo returned a *p* value identical to those returned by TestSym and IQ-TREE.

**Table 1:** Probabilities of obtaining the site-pattern distribution in **N**(*t*_2_) by chance, assuming symmetry, marginal symmetry, and internal symmetry of the evolutionary processes. The probabilities were obtained using Homo v2.0, TestSym (Ababneh et al. 2006b) and IQ-TREE (Naser-Khdour et al. 2020).

| Matched-pairs test of | Homo v2.0 | TestSym v1.0 | IQ-TREE v1.7 |
| --- | --- | --- | --- |
| Symmetry | 0.0213 | 0.0213 | 0.0213 |
| Marginal symmetry | — | 0.1836 | 0.1836 |
| Internal symmetry | — | 0.0183 | 0.0183 |

Foster’s (2004) test is very similar to Stuart’s (1955) matched-pairs test of marginal symmetry, so results obtained from the former test should be compared to those obtained from the latter. Assuming that evolution occurred under the GTR model, Foster’s (2004) test returned a probability of 0.150 and, if it had occurred under the F81 model, 0.157. In summary, Foster’s (2004) test returned lower probabilities than that from Stuart’s (1955) matched-pairs test of marginal symmetry (Table 1), most likely because the two tests used different approaches to assess the same null hypothesis.

Next, we compared the times taken by Homo v2.0 and Homo v1.4 to complete an analysis of the same data. To do so, we analysed an amino-acid alignment from Butler et al. (2009). These data—18 sequences and 412,814 sites—were analysed on a MacBook Air (Processor name: Intel Core i5; Processor speed: 1.6 GHz). Homo 2.0 completed the survey in 0.43 s while Homo 1.4 completed it in 143.26 s; that is a 341-fold speedup. When Homo v2.0 was used in b mode, the essential output was returned in 0.317 s. In conclusion, Homo v2.0 is well-tuned for large phylogenomic data sets.

## Biological Example

To illustrate the insights that may be gained by using the software presented in this paper, we surveyed an alignment of amino acids from Butler et al. (2009). The data matrix is the one used in the previous section.

### The Survey

The PP plot in Figure 6a reveals that this data set is unlikely to have evolved under homogeneous conditions, but a single dot at the righthand side of the plot suggests that at least one pair of sequences have evolved under similar conditions. The heat map in Figure 6b shows that these two sequences come from *Saccharomyces cereviciae* and *S. paradoxus*. The summary statistics for the 153 (non-independent) comparisons show that the smallest *p*-value was 0.0, and that 99.3% of the *p*-values are below the 5% family-wise error rate. In summary, we conclude that the alignment has a strong compositional signal and that only two of the 18 sequences appear to have evolved under the same conditions. Compositional heterogeneity is clearly a pronounced feature of these data, so it would be wise to consider this feature carefully when analysing the data phylogenetically. We note that the large number of sites here can produce very small *p*-values corresponding to small deviations from homogeneity.

### The Impact

An obvious question arising from this discovery is whether the compositional signal is phylogenetic (i.e., whether it, on its own, is able to produce what essentially looks like a phylogenetic tree). To address this question, we analysed the data using the network- and tree-based methods described above.

Figure 7 depicts the compositional network inferred from the **D**_*CFS*_ matrix derived from the multiple sequence alignment of amino acids published by Butler et al. (2009). The network is highly complex and treelike, with several internal splits many times longer than the alternative splits. This feature implies that the phylogenetic tree reported by Butler et al. (2009) may be affected by a strong and complex compositional signal.

To determine whether this is the case, we compared the tree published by Butler et al. (2009) (Fig. 8a) to the compositional tree inferred from **D**_*CFS*_ (Fig. 8b). The important thing to observe here is that five of the internal edges in the two trees are identical. There is no reason to expect the two trees to be more similar or dissimilar to each other than any pair of random trees, so there may be reason to question the accuracy of the phylogenetic tree inferred by Butler et al. (2009). To ascertain whether there is reason for such concern, we compared the two trees statistically.

**Figure 8:**
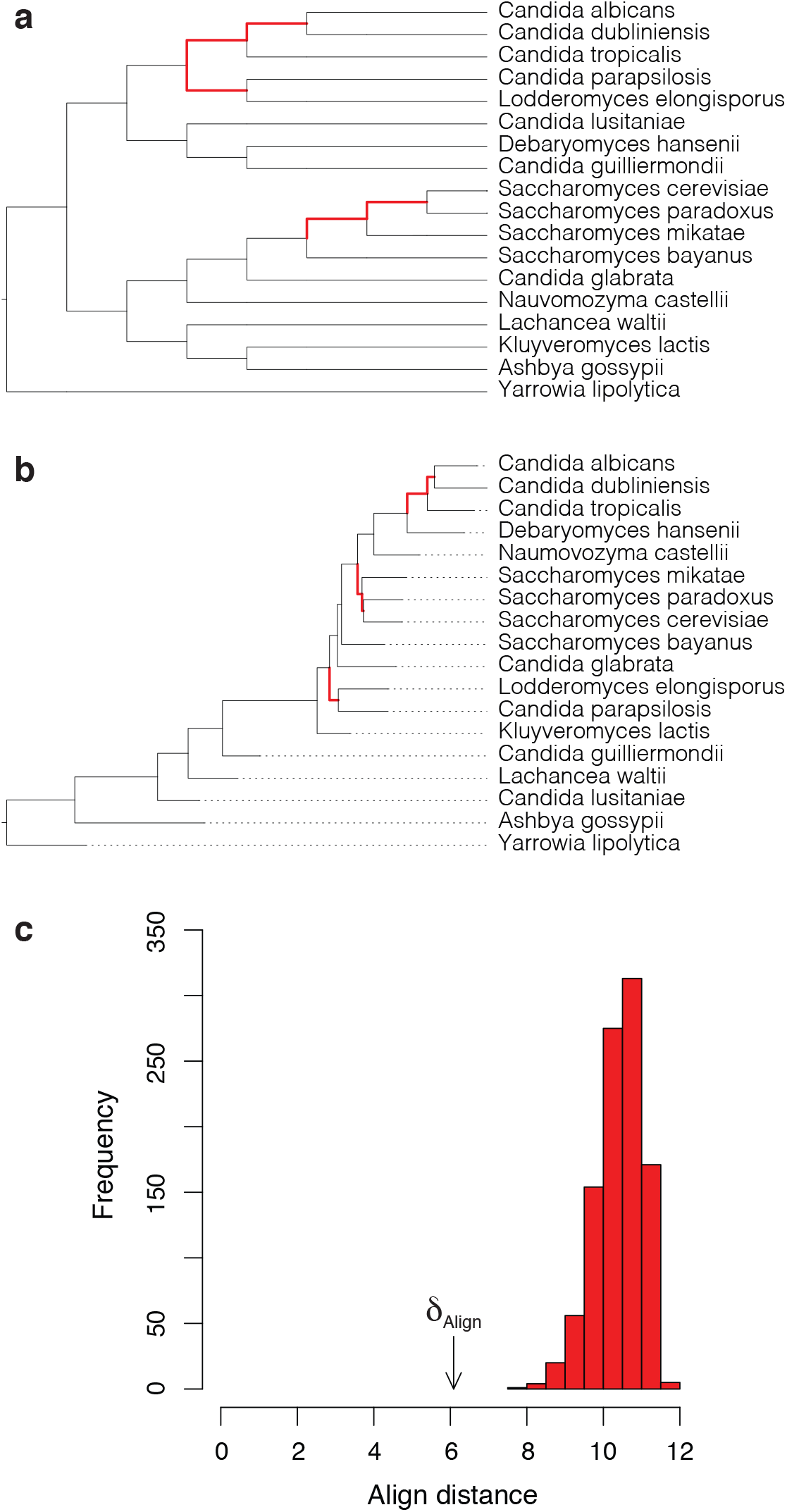
Figure with (**a**) the tree topology inferred by Butler et al. (2009), (**b**) the tree topology inferred from **D**_*CFS*_ using the least-squares distance method implemented in PHYLIP (Felsenstein 2005), and (**c**) a histogram with the align distance between the trees in panels **a** and **b** (arrow) and between 999 randomly-generated pairs of 18-tipped trees (red bars). A similar result was obtained using the quartet distance (Sand et al. 2014). Identical splits in the two trees are highlighted using bold red edges. In Butler et al. (2009), *Lachancea waltii* was called *Kluveromyces waltii* and *Naumovozyma castelliii* was called *S. castellii*.

In practice, we computed *δ*_*align*_ for the two trees in Figure 8 as well as 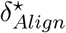 for 999 pairs of randomly-generated 18-tipped trees. The latter estimates were needed to generate the null distribution. Figure 8c shows that the *δ*_*Align*_ value for the two trees lies well below the distribution of 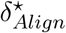 values for the randomly-generated trees, implying that the trees are significantly more alike than random trees are (two-tailed test, *p* < 0.002). Therefore, we may now conclude that the tree topology published by Butler et al. (2009) is affected by the presence of a compositional signal in the alignment of amino acids. In other words, the tree in Figure 1 of Butler et al. (2009) may not reflect the evolution of these 18 species.

## Availability

Homo v2.0 is available from http://www.github.com/lsjermiin/Homo.v2.0/.

HomoHeatMapper is available from http://www.github.com/lsjermiin/HomoHeatMapper/.

RandTree v1.0 is available from http://www.github.com/lsjermiin/RandTree.v1.0/.

## Acknowledgements

We are grateful to Mary Kuhner for processing the data depicted in Figure 8c and to Kenneth Wolfe for constructive feedback.

